# Cell wall composition drives off-target sensitivity to indaziflam

**DOI:** 10.64898/2026.11.10.765741

**Authors:** Mohit Mahey, Manjot Kaur Sidhu, Timothy D. Miles, Peter K. Lundquist, Eric L. Patterson

## Abstract

Indaziflam is a potent alkylazine herbicide classified as a cellulose biosynthesis inhibitor (CBI) with broad-spectrum pre-emergence activity and long soil persistence. While its efficacy in commercial weed control is established, its off-target ecological footprint across taxa with non- cellulose cell wall compositions remains poorly defined. We evaluated the potential off-target biological impacts of indaziflam across organisms representing distinct cell wall architectures, including the yeast *Saccharomyces cerevisiae* (chitin/glucan), the unicellular alga *Chlamydomonas reinhardtii* (glycoprotein), and the early-diverging bryophyte *Marchantia polymorpha* (cellulose/hemicellulose/pectin). Indaziflam did not inhibit the growth of non-target organisms lacking plant-type cellulose cell walls. In *S. cerevisiae*, treatment unexpectedly accelerated growth rates to reach ED_50_ (12.95 h at 3200 pM vs. 16.23 h in control). Similarly, *C. reinhardtii* showed no growth suppression across concentration ranges up to 100 nM. Conversely, the non-vascular land plant *M. polymorpha* exhibited marked, dose-dependent sensitivity. In vitro, thallus surface area was significantly suppressed at concentrations above 1600 pM (*p* < 0.05). In greenhouse trials, pre-emergence application of indaziflam at recommended (1X) and elevated (2X, 3X) field rates provided up to 100% control of liverwort establishment and biomass accumulation over an 8-week evaluation period. Off-target activity of indaziflam is strictly confined to land plants (Embryophytes) possessing conserved cellulose-containing cell walls, with no adverse growth effects observed in organisms featuring chitin- or glycoprotein-based wall structures. These findings clarify the selective environmental toxicity profile of indaziflam, demonstrating that its biological activity aligns closely with its CBI classification without causing widespread toxicity across non-plant soil organisms.

## INTRODUCTION

Herbicides are the primary tools used for weed control in modern commercial agriculture(Norsworthy et al. 2012) forming a major segment of an estimated US$85 billion crop protection market. Given the expansive scale of modern agriculture, chemical herbicides remain the most effective, efficient and economical form of weed control, accounting for US$11.44 billion of sales in 2025, with projected growth to US$14.63 billion by the year 2030(Green 2014, 2026). Herbicide use continues to rise in both developed and developing regions, driven largely by shifting agricultural practices. In particular, conservation tillage and no-till systems that are designed to conserve soil, water and energy, further increase reliance on chemical herbicides for sufficient weed control (Gianessi 2013). Concurrently, the rapid evolution of herbicide-resistant weeds has exacerbated this reliance as growers frequently resort to higher application rates, tank mixing, and more frequent applications(Green 2014). Together, these drivers underscore the urgent need for new herbicide modalities with distinct target sites.

Herbicides are chemicals designed to eliminate or suppress undesirable vegetation(US EPA 2015). Despite their target specificity, numerous off-target effects have been documented. Off-target effects can include injuries or changes in non-target crops, weeds, soil microbes, fungi and other non-target organisms. For example, glyphosate, a widely-used broad spectrum herbicide has been reported to reduce the spore viability and root colonization of arbuscular mycorrhizal fungi even at doses below recommended application rates(2013). Glyphosate has also been shown to disrupt interactions between earthworms and symbiotic mycorrhizal fungi, leading to disruption of the soil ecosystems(Zaller et al. 2014). Further, four species of entomopathogenic fungi *B. bassiana, M. anisopliae, N. rileyi* and *N. floridana exhibit* marked susceptibility to field-relevant concentrations of glyphosate(Morjan et al. 2002). Similarly, paraquat impairs growth of *Trichoderma viride*(Ghannoum et al. 1989); a particularly concerning development as *T. viride* provides important antagonism against several plant pathogens such as *Armillaria mellea, Fomes annosus, Rhizoctonia solani* and *Sclerotium rolfsii*(Wilkinson and Lucas 1969).

Indaziflam is an alkylazine herbicide registered by Bayer Agriscience in 2010 belonging the cell wall biosynthesis inhibitor mode of action (HRAC Group 29)(Myers et al. 2009). It is the active ingredient of commercial formulations such as Alion™ and Specticle™. Indaziflam’s high efficacy, broad-spectrum control of both monocots and dicots, and long residual activity in the topsoil make it ideal for pre-emergence weed control in orchards, rangelands, ornamentals, and turfgrass (Kaapro and Hall 2012). With soil half-life of 150 days, indaziflam (Sebastian et al. 2016, 2017) provides long-lasting (up to 3 years) and effective weed control at low application rates (25-150 g a.i. ha^-1^ hectare).

Although indaziflam is classified as a cellulose biosynthesis inhibitor (CBI), its target site and mode of action are incompletely characterized. To date, there are no reports of direct interaction and/or binding of indaziflam with the cellulose synthesis complex (CSC) nor has cross-resistance between mutants resistant to known CBIs such as isoxaben been demonstrated(Brabham et al. 2014). Thus, it remains unclear whether indaziflam specifically targets plant-type cell walls composed of cellulose, or if it can also inhibit chemically distinct cell walls found in other organisms.

Cell wall composition differs substantially across biological kingdoms(Panstruga et al. 2023). In contrast to plant cell walls, comprised primarily of cellulose and hemicellulose, fungal cell walls consist primarily of chitin and glucan networks with their core scaffold features being covalently linked, branched *β*-1,3-glucans interconnected with chitin via hydrogen bonds. Although subtle variations exist among fungal taxa, this basic architecture is conserved. Compared to plant cell walls, fungal cell walls are more rigid due to extensive inter-chain hydrogen bonding among *N*- acetylglucosamine residues in chitin(Gow et al. 2017; Panstruga et al. 2023). Furthermore, *Chlamydomonas reinhardtii*, as a freshwater microalga, possesses atypical cell wall composed of five dense glycoprotein-rich layers around a water-insoluble central core, lacking the cellulose found in most other microalgae(Poulhazan et al. 2024). Meanwhile, liverworts (division *Marchantiophyta*), which represent one of the earliest diverging lineages of land plants, possess cell walls composed of cellulose, hemicelluloses, and pectins. This makes their cell walls structurally homologous to those of vascular land plants(Pfeifer et al. 2022).

In this study, we evaluated the potential off-target biological impacts of indaziflam across organisms spanning distinct cell wall compositions. Given its extensive use and long soil persistence, we conducted dose–response assays across a panel of organisms - ranging from fungi and oomycetes to the unicellular alga *C. reinhardtii* and the bryophyte *Marchantia polymorpha*. We hypothesized that indaziflam-mediated inhibition would be restricted exclusively to organisms possessing cellulose-based cell walls. Our results indicate that indaziflam’s off-target effects are constricted to cellulose-containing land plants, consistent with their CBI classification.

## MATERIALS AND METHODS

### *Growth of* Saccharomyces cerevisiae

*Saccharomyces cerevisiae* (strain YMK839) was grown overnight in yeast extract peptone dextrose (YPD) medium at 30°C. Cultures were diluted to an optical density at 600 nm (OD_600_) of 0.2, pelleted by centrifugation at 14,000 rpm for 1 min, and resuspended in 200 µL sterile water. A dose-response time-course assay was additionally performed. Yeast was inoculated in 100 mL YPD medium spiked with indaziflam at 100, 200, 400, 800, and 1600 picomolar concentrations and OD_600_ values were recorded every 3 hours over 24 hours using a Genesys 50 spectrophotometer (Thermo Fisher Scientific Inc., MA, USA) using three independent biological replicates.

### *Growth of* Chlamydomonas reinhardtii

The unicellular green alga *Chlamydomonas reinhardtii* (provided by Dr. Christoph Benning, Michigan State University) was cultured in Tris-acetate-phosphate (TAP) medium (Harris, 2009a; Sigma-Aldrich, St. Louis, MO, USA) under 12-hour light/dark cycle at 80 µmol photons m^−2^s^−1^and at 25 °C with shaking at 80 rotations per minute (RPM). Indaziflam was added to cultures at final concentrations of 1.6, 6.4, 25.6, 50, 100, and 500 nanomolar. Cell growth was monitored by measuring OD_600_ using a NanoDrop One spectrophotometer (Thermo Fisher Scientific Inc., MA, USA) at 24-hour intervals for 4 days after treatment.

### Growth of Liverwort on Plates

The Bryophyte, *Marchantiophyta* or liverwort (provided by Dr. Debalina Saha, Michigan State University) was propagated using the gemma produced from a single plant. The gemma was placed on square (10 cm × 10 cm) Petri plates containing 1% agar (Thermo Fisher Scientific Inc., MA, USA) supplemented with half-strength Murashige & Skoog (M519) basal medium. Plates were grown in a growth chamber maintained at 25°C during a 16-hour photoperiod and 21°C during an 8-hour dark period for two weeks. Plates were spiked with filter-sterilized indaziflam at 100, 200, 400, 800, 1600, and 3200 picomolar. The area of the liverwort was measured using FIJI software(Schindelin et al. 2012).

### Pre-Emergence Indaziflam Test on Liverworts

The experiments were conducted in a greenhouse at Michigan State University (MSU) Horticulture Teaching and Research Center. The temperature inside the greenhouse was controlled, with a minimum temperature of 21°C, maximum temperature of 26.6 °C and average temperature of 23.8 °C. The average air temperature was 16°C, air humidity 70%, wind speed 1 mph and precipitation 0 inches. All the sprays were done in an open hoop house. A carbon dioxide (CO2) backpack sprayer (custom built by Bellspray R&D sprayer Inc., Opelousas, LA) calibrated to deliver 252.55 liters/hectare using an 8004 flat-fan nozzle (TeeJet Technologies, Wheaton, IL) at a pressure of 206.843 kilopascals was used for spraying.

In the hoop house, indaziflam was sprayed over the top of container substrate at the rate of 0.55 liters per acre as per the manufacturer’s recommendation (1X). In addition, 2X and 3X application rates were also applied. The next day, gemmae of common liverwort (Marchantia polymorpha) were applied. Gemmae were collected by scraping gemmae cups of vigorous liverwort stock plants and releasing the gemmae into a 250 mL bowl of tap water to separate from their clumps. Approximately 5 mL of water from the bowl, containing approximately 20-25 gemmae, was applied to the surface of each container. The bi-weekly application of gemmae was continued until the end of the experiment. For data collection, visual estimation of percent control of liverwort was done bi-weekly for 8 weeks. The fresh weight of liverwort was determined for each pot at the end of the experiment. The experiment was conducted in a completely randomized design including three rates of herbicide application and an untreated control. There were six single-pot replications per treatment.

### Statistical Analysis and Dose-Response Curves

The growth data from *Chlamydomonas reinhardtii* and *Saccharomyces cerevisiae* was analyzed using the drc (v3.0-1) R package(Ritz et al. 2015). Growth data was analyzed using a three- parameter log-logistic dose-response model to capture the various growth stages, namely lag, log and stationary phase. Model performance was assessed using, Akaike Information Criterion (AIC), Bayesian Information Criterion (BIC), and Root Mean Squared Error (RMSE) implemented in performance (v0.15.0) R package (**Supplementary Table 1**). From the fitted models, effective doses (ED_50_) were estimated relative to the maximal response, providing a standardized measure of the time required for each treatment to achieve defined inhibition levels. For analysis of liverwort and *Phytophthora capsici*, one-way ANOVA analyses were performed using GraphPad Prism (DotMatics, San Diego, USA). All analyses were conducted using R (v4.4.0)(R Core Team 2024) with the tidyverse (v2.0.0)(Wickham et al. 2019) and multicompView (v0.1-10) R packages(Graves et al. 2019).

For pre-emergent treatment of liverwort with indaziflam, the data was analyzed using a generalized linear mixed model in by PROC GLIMMIX in SAS (Ver. 9.4, SAS Institute, Cary, NC) to conduct the two-way Analysis of Variance (ANOVA). When ANOVA results revealed significant effects, mean comparisons for rates of herbicide were performed using Tukey’s Honest Significant Difference (HSD). All the effects were considered significant at alpha=0.05, to separate out the means.

## RESULTS

### Effect of Indaziflam Treatment on Saccharomyces cerevisiae Growth

We first investigated indaziflam inhibition of the single-celled fungal species, *Saccharomyces cerevisiae*, which has a chitin-based cell wall. Time-course growth of *S. cerevisiae* exposure to varying indaziflam concentrations is shown in **Figure 1**. The three-parameter log-logistic model fit the OD_600_ data well (AIC = -177.419, BIC of -106.101, RMSE of 0.135; **Supplemental Table S1**). A lack-of-fit test showed no evidence of model misfit relative to a saturated ANOVA model (F = 0.651, p = 0.945).

**Figure 1.**
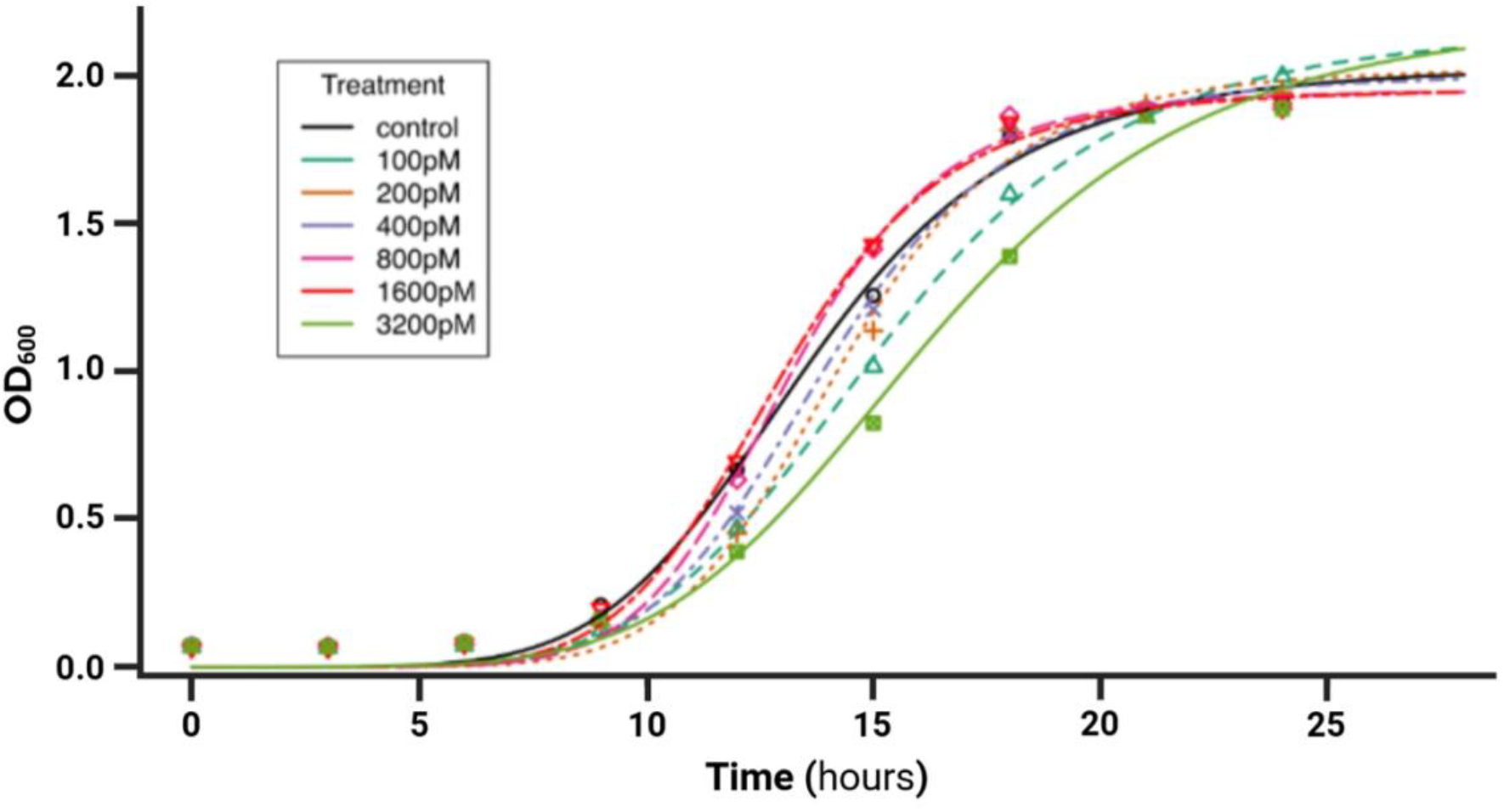
Dose-response growth curves for *Saccharomyces cerevisiae* treated with indaziflam. *S. cerevisiae* was grown in liquid culture spiked with indaziflam at the specified concentrations. OD_600_ was recorded every three hours over the growth cycle, capturing lag, exponential and stationary growth phases. Data points illustrate measured OD_600_ values at given time points. Colored lines represent three-parameter log-logistic (LL.3) model best-fits by treatment. Tested concentrations range from 0 - 3200 pM final concentration.

Effective dose estimates were calculated at 50% maximum growth (ED_50_). For the untreated control, ED_50_ was 16.23 hours. *S. cerevisiae* exhibited a dose-dependent decrease in the time required to reach 50% growth. At 3200 pM the ED_50_ was reduced to 12.95 hours compared to an ED_50_ of 16.23 hours in control media, indicative of a faster growth rate with increasing indaziflam concentrations (**Table 1**).

**Table 1.** Estimated effective doses (ED_50_) for *Saccharomyces cerevisiae* under indaziflam treatments.

| Indaziflam concentration (pM) | Dose (ppm) | Estimated ED<sub>50</sub> (hours)<sup>*†*</sup> |
|---|---|---|
| Control | – | 16.2 ± 0.8 |
| 100 | 3.01×10<sup>−5</sup> | 13.5 ± 0.4 |
| 200 | 6.03×10<sup>−5</sup> | 15.2 ± 0.6 |
| 400 | 1.21×10<sup>−4</sup> | 14.3 ± 0.3 |
| 800 | 2.41×10<sup>−4</sup> | 13.9 ± 0.3 |
| 1600 | 4.82×10<sup>−4</sup> | 13.1 ± 0.3 |
| 3200 | 9.64×10<sup>−4</sup> | 13.0 ± 0.3 |

### Effects of Indaziflam Treatment on Chlamydomonas reinhardtii Growth

Next, we investigated indaziflam inhibition of the photosynthetic single-celled algal species, *Chlamydomonas reinhardtii*, with a glycoprotein cell wall composition. The three-parameter log- logistic model was used to model *C. reinhardtii* growth measured at OD_600_ (**Figure 2**), with good model fit (AIC = 55.049, BIC = 98.305, RMSE = 0.272; **Supplemental Table S1**). At the highest indaziflam concentration of 100 nM *C. reinhardtii* reached ED_50_ in 2.0 days, compared to the untreated control conditions where it reached ED_50_ in 1.7 days, no statistically significant difference was observed between these treatment doses (**Table 2**).

**Table 2.** Estimated effective doses (ED_50_) for *Chlamydomonas reinhardtii* under indaziflam treatments.

| Indaziflam concentration (nM) | Dose (ppm) | Estimated ED<sub>50</sub> (days)<sup>*†*</sup> |
|---|---|---|
| Control | – | 1.7 ± 0.6 |
| 1.6 | 4.82×10<sup>−4</sup> | 2.1 ± 1.0 |
| 6.4 | 1.93×10<sup>−3</sup> | 3.1 ± 3.1 |
| 25.6 | 7.71×10<sup>−3</sup> | 2.3 ± 0.9 |
| 50 | 1.51×10<sup>−2</sup> | 2.3 ± 1.0 |
| 100 | 3.01×10<sup>−2</sup> | 2.0 ± 0.7 |

*† Mean ± 1 s.e.m.*

**Figure 2.**
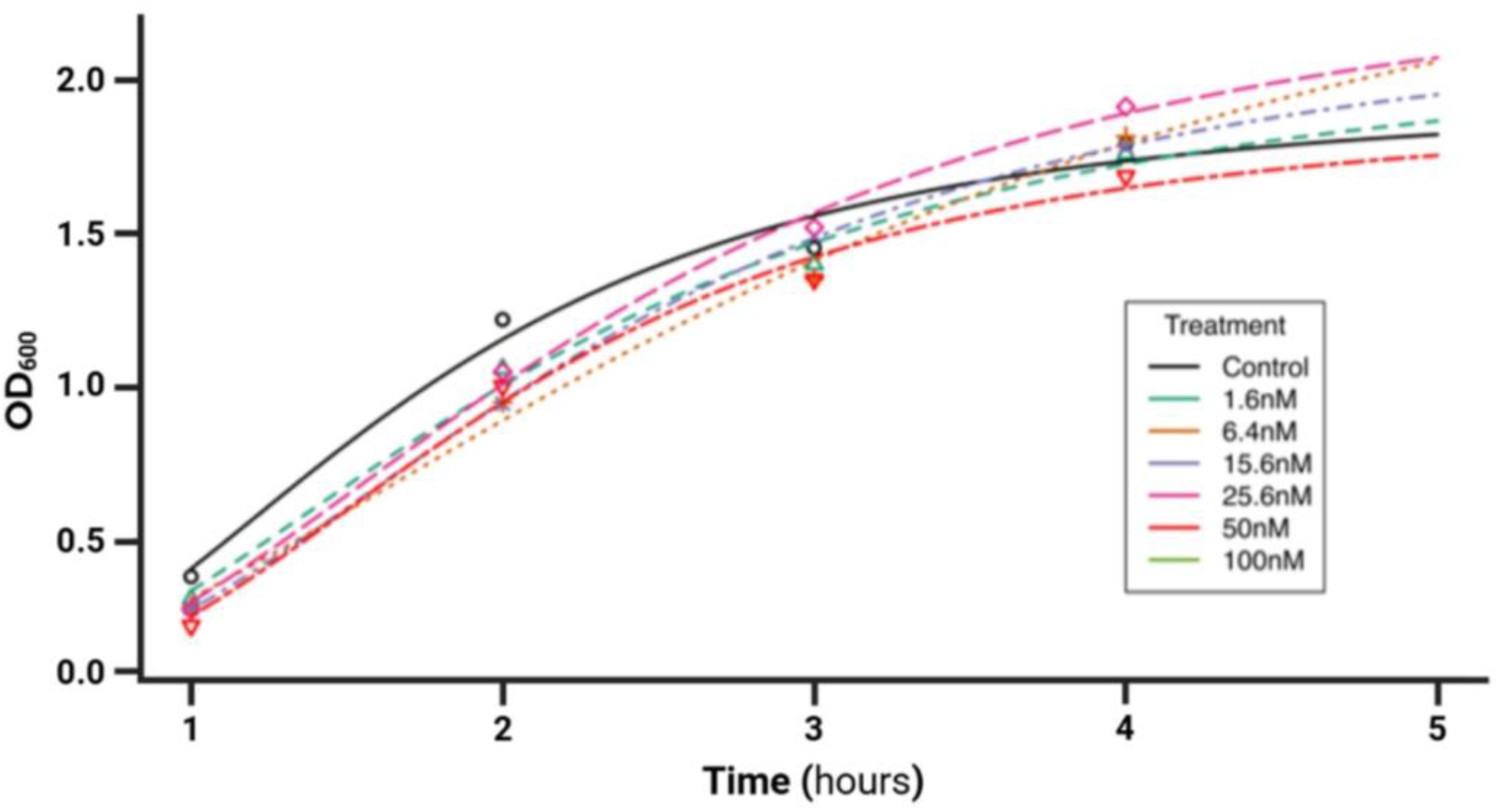
Dose-response growth curves for *Chlamydomonas reinhardtii* treated with indaziflam. *C. reinhardtii* was grown in liquid culture spiked with indaziflam at the specified concentrations. OD_600_ was recorded daily over the growth cycle, capturing population growth trajectories under control (0 mM indaziflam) and treated conditions. Data points illustrate measured OD_600_ values at given time points. Colored lines represent three-parameter log-logistic (LL.3) model best-fits by treatment. Tested concentrations range from 0 - 100 nM final concentration.

### Effects of Indaziflam Treatment on In Vitro Oomycete Growth

Next, we investigated indaziflam inhibition of the the oomycete, *Phytophthora capsica*, with a more plant-like cellulose (β-1,4-glucan) cell wall. Colony sizes were not affected in a dose- dependent manner (**Figure 3**), although colony sizes did fluctuate somewhat across differing indaziflam concentrations. We conclude that *Phytophthora capsica* growth is not sensitive to indaziflam within the concentrations tested.

**Figure 3.**
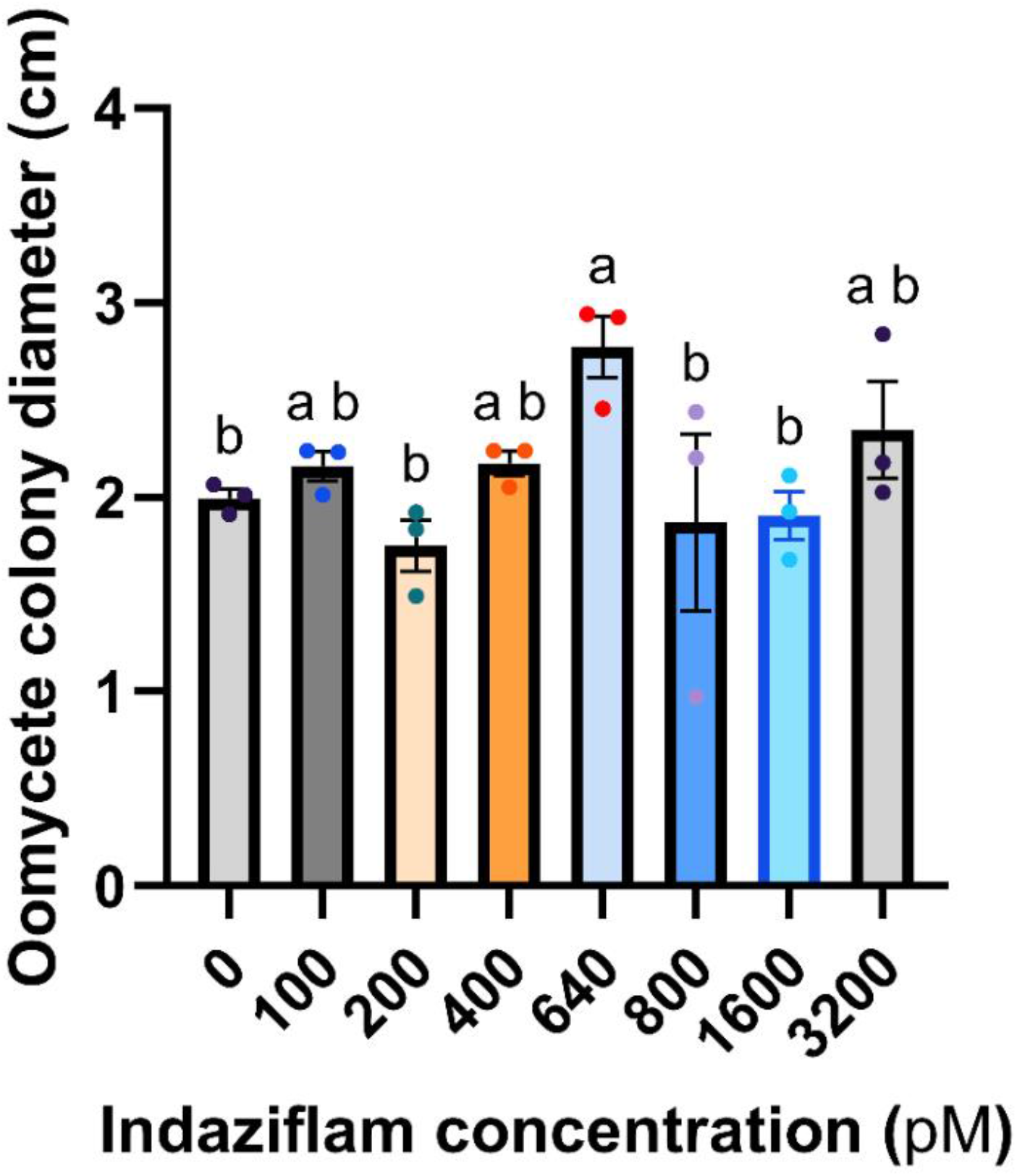
Oomycete growth response to indaziflam treatment. Oomycete (*Phytophthora capsici*) was grown on agar plates spiked with indaziflam at the specified concentrations. Colony diameters were measured using FIJI software and recorded after 14 days of growth on treated agar. Bars represent the mean ± 1 s.e.m; n = 3 biological replicates (individual oomycete colonies). Statistically significant differences between treatments are determined by ordinary one-way ANOVA with Fisher’s LSD test, indicated using single-letter codes.

### Effects of Indaziflam Treatment on In Vitro Liverwort Growth

We then investigated the effect of indaziflam treatment on liverwort, *Marchantia polymorpha*, with its plant-like cellulose/hemicellulose-based cell wall composition. Indaziflam significantly reduced thallus surface area *in vitro* (F = 7.06, *p* = 3.3 × 10^−6^). Thallus growth decreased markedly at higher concentrations, with the greatest suppression observed at 3200 pM (**Figure 4**). Pairwise Tukey HSD comparisons confirmed that 3200 pM caused a strong and statistically significant reduction in liverwort area relative to the control (*p* = 0.00095). Similarly, thallus area at 1600 pM treatment also resulted in significantly lower area compared with 100 pM (*p* = 0.021) and marginally lower area than 800 pM (p = 0.057). Lower concentrations (100 - 800 pM) did not differ significantly from each other or from the control. Thus, liverwort growth remained stable across relatively low doses (< 1600 pM) but decreased sharply above 1600 pM.

**Figure 4.**
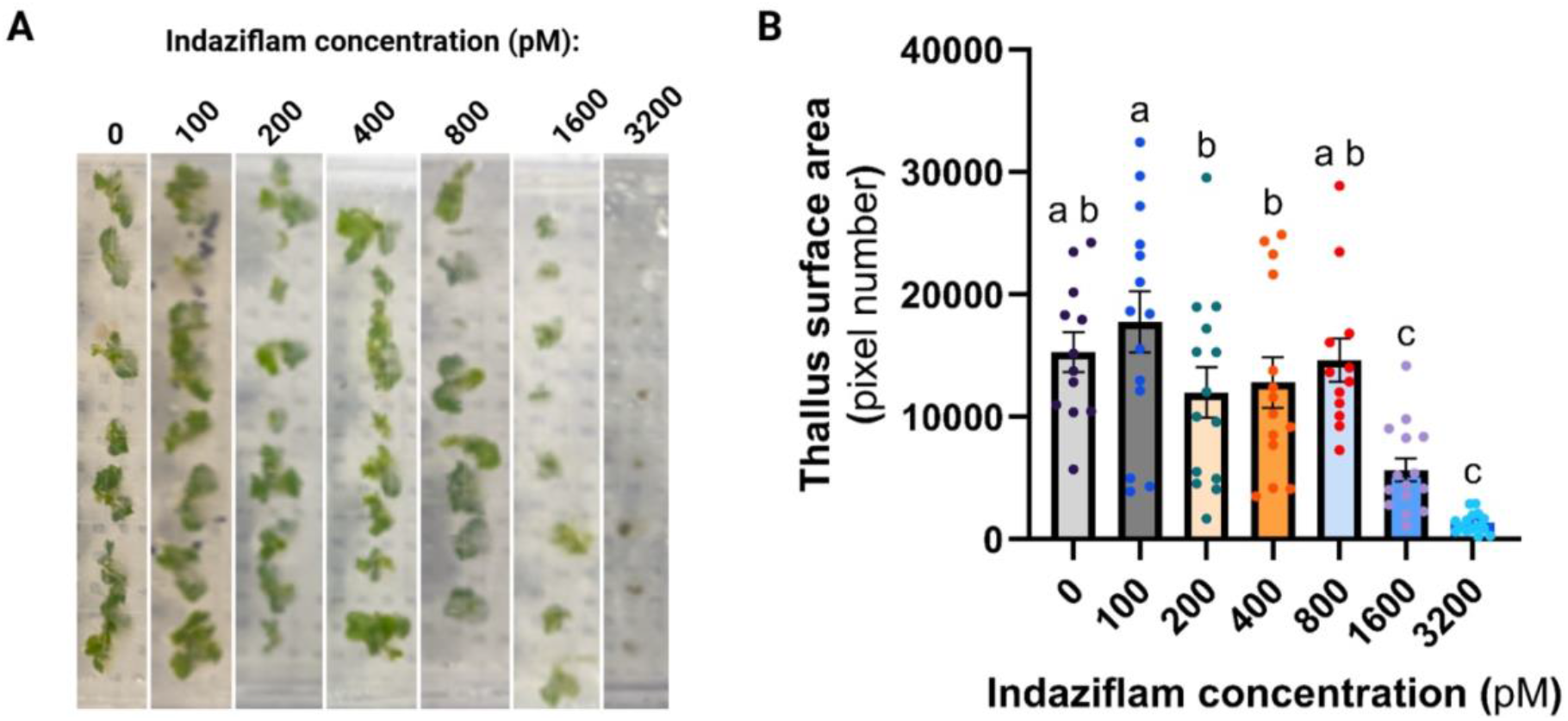
Liverwort growth response to indaziflam treatment. Liverwort (*Marchantia polymorpha*) was grown on agar plates spiked with indaziflam at the specified concentrations.(A) Images of liverwort thalli grown on agar spiked with indaziflam B) Surface area of liverwort thallus was measured using FIJI software as total pixel number and recorded after 14 days of growth on treated agar. Bars represent the mean ± 1 s.e.m; n = 12 - 15 biological replicates (individual liverwort thalli). Statistically significant differences between treatments are determined by ordinary one-way ANOVA with Fisher’s LSD test, indicated using single-letter codes.

### Pre-Emergence Control of Liverwort in Greenhouse Trials

To further investigate the effect of indaziflam on liverwort growth, we tested pre-emergence applications of indaziflam in greenhouse trials. Pre-emergence applications of indaziflam across all tested application rates (1X, 2X and 3X) provided significant control of *M. polymorpha* throughout the 8-week evaluation period (p<0.05) (**Table 3**). At all bi-weekly time points, liverwort coverage in indaziflam-treated pots was significantly lower than in untreated controls, though differences among application rates were minor.

**Table 3.** Liverwort growth characteristics in presence of indaziflam applied pre-emergence.

| Application Rate | Liverwort coverage (%)<sup>†</sup> |  |  |  | Fresh biomass (g)<sup>§</sup> |
|---|---|---|---|---|---|
|  | 2 WAT<sup>‡</sup> | 4 WAT<sup>‡</sup> | 6 WAT<sup>‡</sup> | 8 WAT<sup>‡</sup> |  |
| Control | 10.5<sup>a, ¶</sup> | 51.7<sup>a</sup> | 78.5<sup>a</sup> | 91.2<sup>a</sup> | 45.3<sup>a</sup> |
| 1X | 2.7<sup>b</sup> | 16.7<sup>b</sup> | 29.0<sup>b</sup> | 31.2<sup>b</sup> | 12.0<sup>b</sup> |
| 2X | 0.0<sup>b</sup> | 0.0<sup>b</sup> | 0.0<sup>b</sup> | 0.0<sup>b</sup> | 0.0<sup>b</sup> |
| 3X | 0.0<sup>b</sup> | 0.0<sup>b</sup> | 0.0<sup>b</sup> | 0.0<sup>b</sup> | 0.0<sup>b</sup> |
| p value | <0.0001 | <0.0001 | <0.0001 | <0.0001 | <0.0001 |

*† Visual estimation of liverwort thallus coverage (%) on the top of the substrate in containers treated with indaziflam applied pre-emergence*
*‡ WAT is Weeks After Treatment, i.e. 2, 4, 6, or 8 weeks after indaziflam application*
*§ Fresh biomass was measured at the conclusion of the experiment (8 weeks after treatment)*
*¶ Percentages followed by the same letter are not statistically significantly different within a column; mean comparisons were performed using Tukey's Honest Significant Difference (HSD) at p = 0.05.*

Across treatment rates, liverwort surface coverage ranged from 0-2.7% at 2 WAT, 0-16.7% at 4 WAT, 0-29% at 6 WAT and 0-31.2% at 8 WAT, compared to 10.5%, 51.7%, 78.5% and 91.2% in untreated control pots at the same timepoints. Meanwhile, the 2X and 3X rates achieved essentially complete inhibition of liverwort growth throughout the period of study (**Table 3**). At 8 WAT, mean fresh thallus biomass was 12.03 g in 1X treatment pots, and 0 g (no living tissue) in 2X and 3X treatment pots, all significantly lower than that recorded for control (45.3 g).

## DISCUSSION

This study evaluated the off-target biological activity of indaziflam across non-target taxa selected for their distinct cell wall structures. Indaziflam has a long residual in the soil, with notable weed control up to three years, as well as broad-spectrum activity at relatively low application rates (25- 150 g a.i. ha^-1^)(Kaapro and Hall 2012). As herbicide resistance spreads globally, growers rely increasingly on newer chemical modes of action. Given their widespread environmental release, understanding the off-target ecological footprint of these chemistries is essential. Previous studies have reported that indaziflam alters the composition of the soil microbiomes, while others report localized increases in microbial biomass, suggesting divergent interactions of indaziflam with micro-organisms as either potent inhibitors or supplemental carbon source(Bradbury et al. 2024)^-^ (Torres et al. 2018).

In our experiments, indaziflam did not cause inhibition in the fungal species tested. Because indaziflam is classified as a cellulose biosynthesis inhibitor (CBI), this lack of fungal toxicity is consistent with the chitin-based architecture of fungal cell walls(Ruiz-Herrera and Ortiz- Castellanos 2010). Similarly, radial growth of the oomycete *Phytophthora capsici* was largely unaffected across all tested indaziflam concentrations (**Figure 3**). Although oomycete cell walls contain β-glucans similar to those in plant walls, their enzymatic assembly machinery diverges sufficiently from plant cellulose synthase complexes to prevent indaziflam inhibition(Hardham 2007). Likewise, the green alga *Chlamydomonas reinhardtii* displayed no evident growth inhibition (**Figure 2**), which aligns with the unique composition of its cell wall, which is made up of five layers of glycoproteins.

Conversely, *Saccharomyces cerevisiae*, exhibited faster growth under higher indaziflam treatment (**Figure 1**), potentially reflecting a hormetic stress response or low-level metabolic utilization of the compound. In striking contrast to the other organisms, the liverwort *Marchantia polymorpha* exhibited clear, dose-dependent sensitivity both in vitro and in greenhouse conditions (**Figure 4**). Liverworts are non-vascular plants (bryophytes) representing one of the earliest lineages of land plants, sharing conserved cell wall synthesis machinery with vascular plants. Their strong susceptibility demonstrates that indaziflam’s inhibitory activity extends across the *Embryophyte* clade to non-vascular land plants, while leaving non-plant and non-cellulose-walled organisms unharmed.

Collectively, these findings demonstrate that indaziflam’s off-target activity is confined to land plants (i.e. Embryophytes) that share conserved cellulose synthesis pathways. To further trace the evolutionary lineage of indaziflam sensitivity, future research should expand testing across additional early-diverging land plants and should include mosses (e.g. *Physcomitrium patens*) and hornworts, as well as charophyte algae, the closest algal ancestors of land plants. Finally, field- based ecotoxicological studies are required to verify whether these laboratory observations translate to complex natural soil ecosystems.

## ACKNOWLEDGEMENTS

The authors wish to thank Dr. Linda Hanson, John Lemas, Dr. Debalina Saha, Dr. Ray Hammerschmidt, Dr. Carrie Hiser, and Hsin Ying Lin (Michigan State University) for assistance in conducting the experiments. Funding was provided by USDA-NIFA Hatch grant to PKL (MICL-02944), Michigan Department of Agriculture and Rural Development (MDARD) Horticulture Fund 2024-25 grant number 240000003331 and Western Michigan Greenhouse Association 2024 grant (to EP and MM).

